# Germline-Restricted Chromosome (GRC) is Widespread among Songbirds

**DOI:** 10.1101/414276

**Authors:** Anna A. Torgasheva, Lyubov P. Malinovskaya, Kira S. Zadesenets, Tatyana V. Karamysheva, Elena A. Kizilova, Inna E. Pristyazhnyuk, Elena P. Shnaider, Valeria A. Volodkina, Alsu F. Saiftdinova, Svetlana A. Galkina, Denis M. Larkin, Nikolai B. Rubtsov, Pavel M. Borodin

## Abstract

The genome of flying birds, the smallest among amniotes, reflects overweight of the extensive DNA loss over the unrestricted proliferation of selfish genetic elements, resulted in a shortage of repeated sequences and lack of B-chromosomes. The only exception of this rule has been described in zebra finch, which possesses a large germ-line restricted chromosome (GRC), transmitted via oocytes, eliminated from male postmeiotic cells and absent in somatic cell. It is considered as a rarity and its origin, content and function remain unclear. We discovered that all songbirds possess GRC: in various size and genetic content it is present in all fifteen songbird species investigated and absent from germ-line genomes of all eight species of other bird orders examined. Our data based on fluorescent *in situ* hybridization of DNA probes derived from GRCs of four different Passeri species and their sequencing indicate that the GRCs show low homology between avian species. They contain fragments of the somatic genomes, which include various unique and repetitive sequences. We propose that the GRC has formed in the common ancestor of the extant songbirds and undergone subsequent divergence. GRC presence in the germ line of every songbird studied indicate that it could contain genetic element(s) indispensable for gametogenesis, which are yet to be discovered.

---

Eukaryotic genomes harbors various selfish genetic elements (transposons, B chromosomes, etc), which enhance their own transmission and might serve as a motors for evolutionary change and innovation^1^. In flying birds, the natural selection led to a reduction of genome size at the expense of transposable elements, introns, constitutive heterochromatin, paralogous genes and other repeated sequences. Resulted genomic compaction provides an economy of bird body mass, improving their metabolic efficiency^2^. An interesting way of resolving a conflict between the body mass and genome size was found in two closely related pet species of Estrildidae birds: zebra and Bengalese finches^3,4^. In all germ line cells, these species contain a large additional acrocentric chromosome, which was absent in somatic cell lines (bone marrow, liver, muscles). In oocytes, this germ-line restricted macrochromosome (GRC) is usually present in two copies as a recombining bivalent. In spermatocytes, one copy of this chromosome forms a round heterochromatic body, which is eliminated from the nucleus during I meiotic division. It has been suggested that GRC might contain multiple copies of genes important for germ cell development and dispensable for soma^3-5^. However, genetic content of GRC, its origin and phylogenetic distribution remains unknown.

Here, using antibodies to the core proteinaceous structure of meiotic chromosomes, the synaptonemal complex (SC), we show the GRC is present in all 15 examined songbird species (13 from this study and two from the previous studies) representing eight families of Passeri (Fig. 1). In eight species, the GRCs were presented by large acrocentric macrochromosomes (macro-GRCs), which were absent from the bone marrow cells (Fig. 2A, B and Supplementary Fig. 1). In oocytes macro-GRC was usually present in two synapsed copies (as a bivalent), which contained one or two terminally located recombination sites visualized by antibodies to MLH1 mismatch repair protein. In the spermatocytes, it usually occurred as a univalent lacking recombination sites and diffusely labeled with centromere antibodies (Fig. 1A and Supplementary Fig. 1). At the end of the male meiotic prophase, the GRC becomes transformed in a dense round body and ejected from the nucleus (Supplementary Fig. 2). A similar meiotic behavior has been described for GRCs in zebra and Bengalese finch^3,4^. In male germline cells of five other species we detected micro-GRC appeared as a univalent without recombination sites surrounded by a cloud of centromere antibodies similar to that described for macro-GRCs. In the oocytes of these species, GRC formed a bivalent indistinguishable from other microchromosomes. We did not observe any phylogenetic clustering for the GRC size. Both macro- and micro-GRCs were present within the families Fringillidae and Hirundinidae (Fig. 1).

**Fig. 1.**
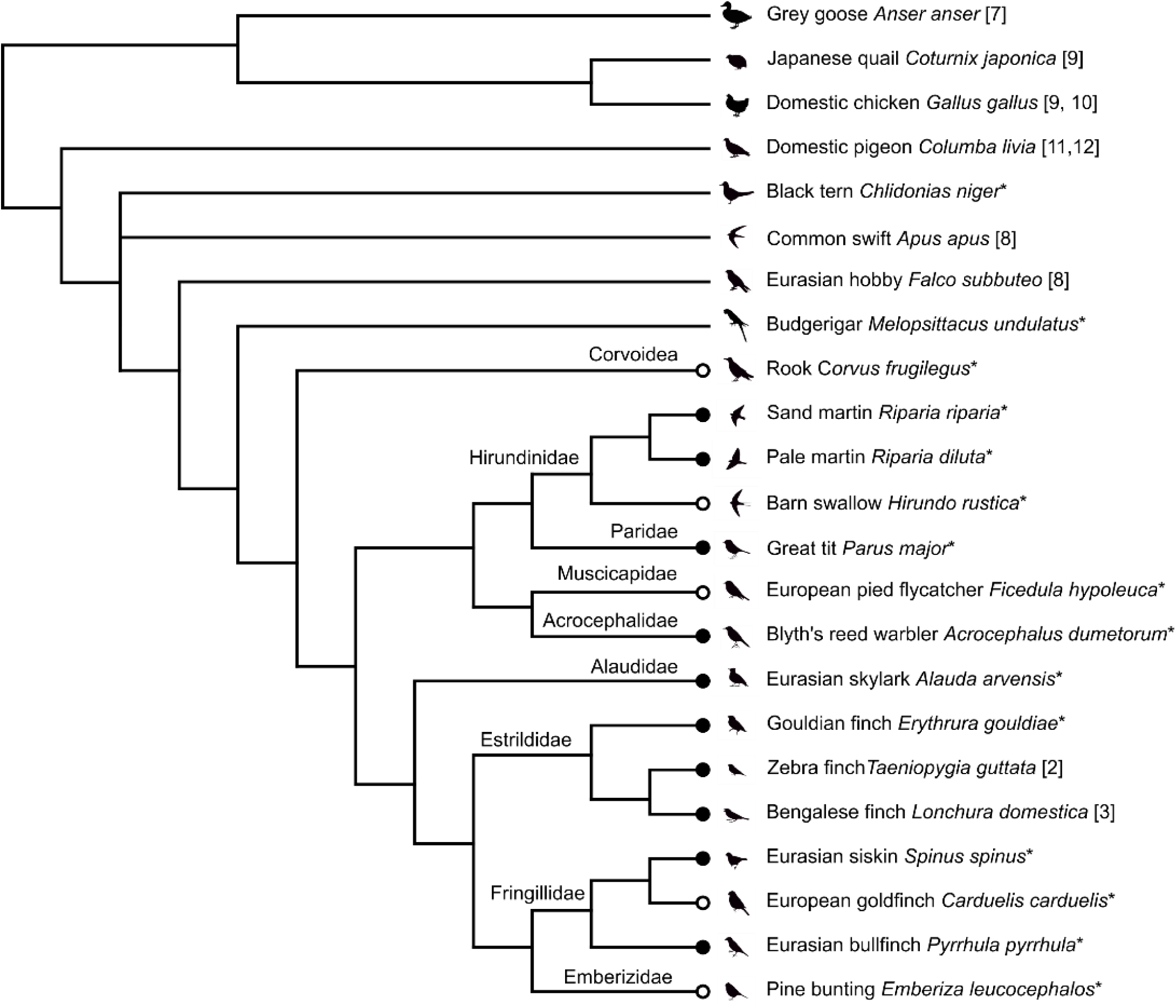
Topology of the bird species examined. Black circles indicate species with macro-GRC, white circles – species with micro-GRC. Numbers after the species names indicate references for SC studies, asterisks are indicative of our data. Consensus topology was based on the cladogram from Reddy et. al^12^. Position of Acrocephalidae and Alaudidae was added according to the TimeTree Database^13^.

**Fig. 2.**
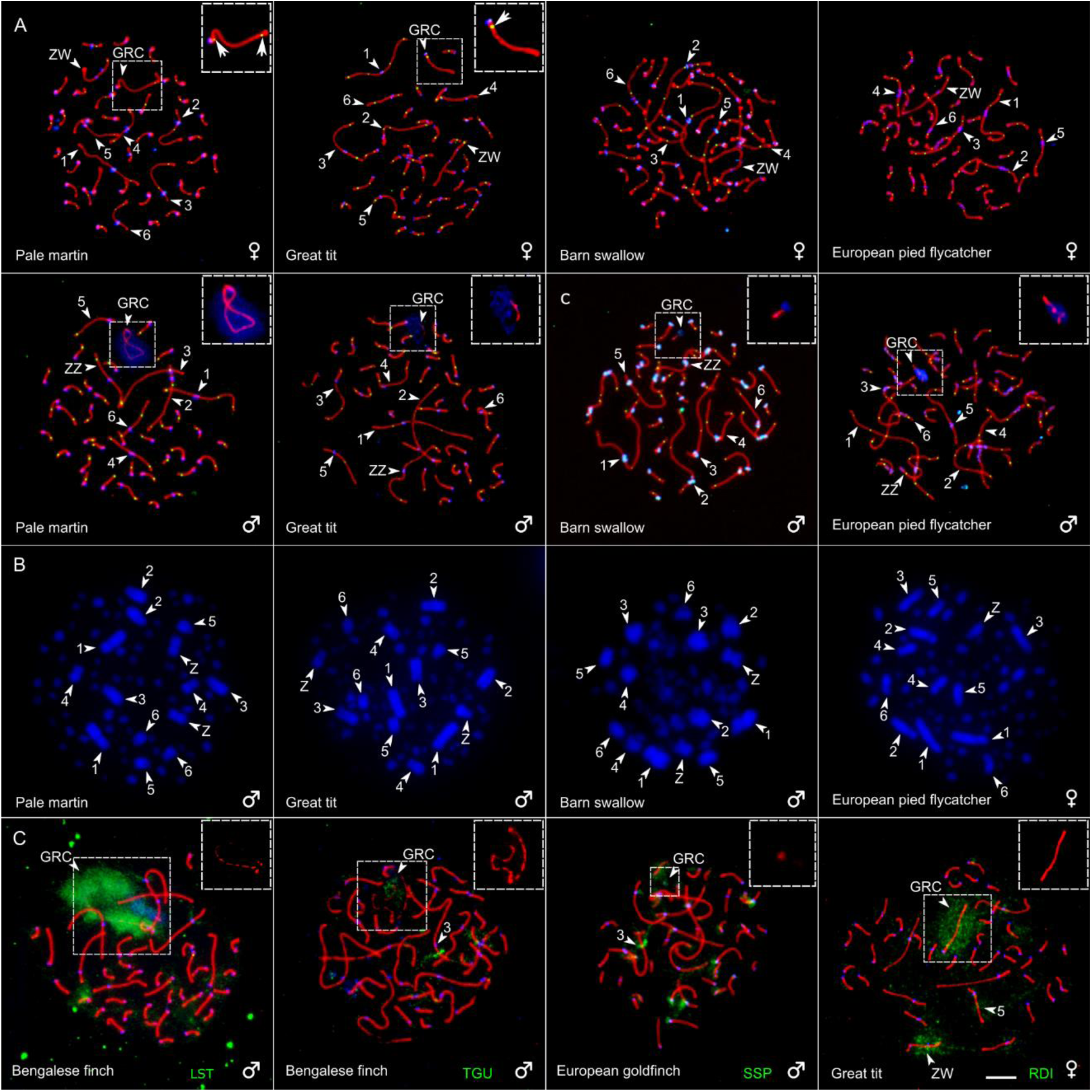
Discovery of GRCs in bird species. (A) SC spreads of four oscine species immunolabelled with antibodies against SYCP3, the main protein of the lateral element of SC (red), centromere proteins (blue) and MLH1, mismatch repair protein marking recombination sites (green). Arrowheads indicate the largest chromosomes ordered according their size ranks, ZZ (identified by its size and arm ratio), ZW (identified by heteromorphic SC and misaligned centromeres), and GRCs. Arrows in the inserts point to MLH1 foci at GRCs. Micro-GRC bivalents in female barn swallow and European pied flycatcher are undistinguishable from microchromosomes of the somatic chromosome set. **(B)** DAPI stained bone marrow cells. **(C)** Reverse and cross-species FISH of GRC DNA probes (green) derived from Bengalese finch (LST), zebra finch (TGU), Eurasian siskin (SSP), and pale martin (RDI) with SC spreads, immunolabelled with antibodies against SYCP3 (red). Centromeres are labeled with antibodies against centromere proteins (blue). Arrowheads indicate GRCs and regions on the somatic chromosome set intensely painted with GRC probes in cross-species FISH. Inserts show GRCs. The Bengalese finch GRC-specific DNA probe produces strong signal on the Bengalese finch GRC and slightly paints some regions of somatic chromosome set. The zebra finch GRC probe paints the distal area of the Bengalese finch GRC and a region of the short arm of SC 3. Eurasian siskin GRC probe paints a micro-GRC of European goldfinch, a region on the long arm of SC 3 and some pericentromeric regions. The pale martin GRC probe produces dispersed signal on the great tit GRC, ZW bivalent and on SC5. Bar – 5 µm.

In any specimen of any examined species, we never detected a single spermatocyte or oocyte without GRC. This indicates that GRC is likely to be an indispensable component of the songbird germline genome (including the rook, the infraorder Corvides). We found no indications of GRC in eight species beyond this Suborder reanalyzing our own data^6,7^ and published SC images^8-11^ (Fig. 1). This might suggest a monophyly of GRC. The estimated time of songbird divergence is 44 MYA (CI: 36 - 50 MYA)^13^. However, as no suboscine species has been examined yet, we cannot exclude a possibility of its appearance in the common ancestor of all Passeriformes about 82 MYA (CI: 75 - 90 MYA)^13^.

To estimate sequence homeology between GRCs of different species and to get insight into their genetic content, we prepared DNA probes of macro-GRCs for four representatives of three families: Estrildidae (zebra and Bengalese finches), Fringillidae (Eurasian siskin), and Hirundinidae (pale martin). We microdissected the round dense bodies (Supplementary Fig. 2) containing GRC from spermatocyte spreads and carried out whole-genome amplification (WGA) of the dissected material. The resulting probes were used for fluorescent *in situ* hybridization (FISH) and NGS sequencing.

Reverse FISH with GRC probes produced strong specific signals on GRCs of each species proving that the round dense bodies are indeed the ejected GRCs (Fig. 1C and Supplementary Fig. 3). In the cross-species FISH experiments, the intensity of specific GRC signals was lower. Importantly, the micro-GRCs were painted with DNA probes derived from macro-GRCs of closely related species. This indicates that GRCs of different species share at least a part of their genetic content. In both reverse and cross species FISH we also detected some signals on somatic chromosomes. Some signals remained visible after repeat suppression with Cot-1 DNA. This indicates that GRCs contain multiple copies of sequences homologous to disperse or/and tandem genomic repeats as well as sequences homeologous to unique regions present in somatic genome.

To identify these sequences we aligned the GRC NGS reads to the repeat-masked zebra finch reference genome (Taeniopygia_guttata-3.2.4^14^) using BLAT^15^ with the 90% identity setting. Genome average coverage estimated in 10 kb windows was 0.15 ± (S.D.) 4.60, 0.12 ± 3.29, 0.03 ± 1.16, and 0.01 ± 0.25 for reads of zebra finch, Bengalese finch, Eurasian siskin, and pale martin GRC libraries, respectively. The coverage was highly uneven. Different species show a homeology to different regions of the reference genome. In total for four GRC libraries, we characterized 27 regions longer that 10 kb, covered by at least 30% of their length and with excess by two standard deviation over the genome average (Table 1). In some regions, where GRC of one species showed a very high coverage, GRCs of other species showed much lower, but still above the average. This might indicate that the unique sequences located in these regions have been copied from the ancestral somatic genome into the ancestral GRC and then diverged in their copy number in the GRCs of different songbirds.

**Table 1.**
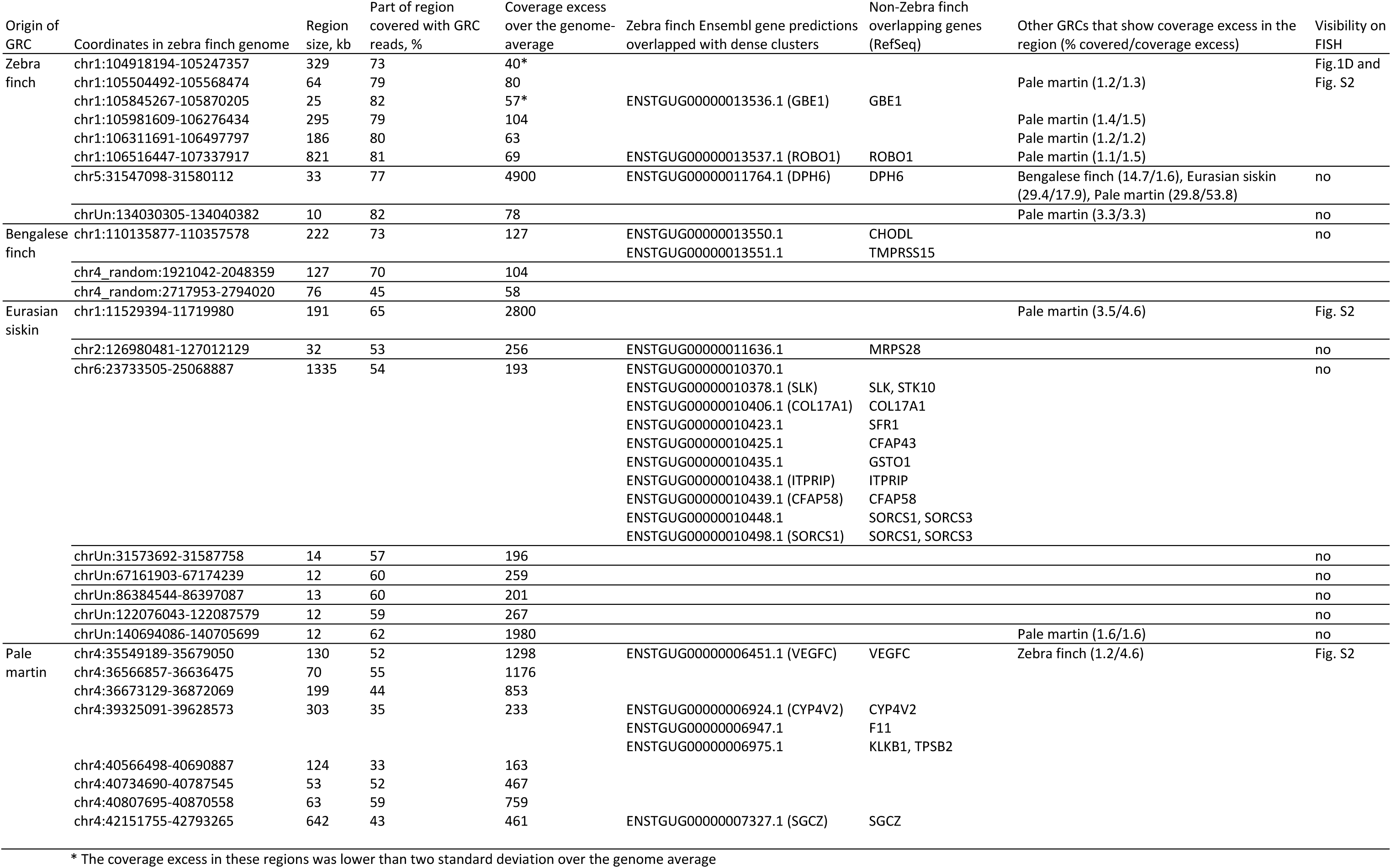
Position of dense clusters of GRC reads aligned to the zebra finch reference genome

The longest regions matched the positions of GRC FISH signals for the corresponding species. Some regions partially overlapped with sequences of zebra finch genes^16^ or sequences homologous to non-zebra finch RefSeq genes^17^ (Table 1). For example, zebra finch GRC probe demonstrated a strong hybridization signal on a short arm of zebra finch SC3 (corresponding to TGU1) and on one of the largest SCs of other species examined (Fig. 1C and Supplementary Fig. 3). In the corresponding region of TGU1, we found a 2.5 Mb long cluster of several regions with ∼70 fold coverage excess (Table 1). This cluster overlapped with two genes: completely with *ROBO1,* a gene involved in vocal learning^18^, and partially with *GBE1*, a gene controlling 1,4-alpha-glucan branching enzyme 1^17^. Homology between the zebra finch GRC and a part of this genomic interval on TGU1 has been detected earlier by the RAPD-PCR technique^19^. Further functional analysis would be required to test whether copies of genes found in the GRC libraries are complete and active or not.

GRCs also contained multiple repeated sequences. We estimated its representation in the GRC reads and in the somatic genome of the zebra finch using RepeatMasker^20^ with the RepBase avian library^21^ (Table 2). RepeatMasker revealed some simple and low complexity repeats. The fraction of transposable elements (TEs) in the GRCs was typical for avian genomes^22^. The majority of them were LTRs and LINEs, while SINEs and DNA TEs were very rare. Overall abundance of LTRs and LINEs and their ratio varied between the GRCs that might reflect different evolutionary trajectories of the GRCs in different species. It has been shown that although activity of TEs in bird genome was rather low and ancient, the species differed for the timing of activity peaks of different TEs. Interestingly, zebra finch genome shows a peak of LTR activity from 5 to 20 MYA^22^. This might be a reason why LTRs are more abundant in zebra finch GRC than in other GRCs. On the other hand, SINEs are extremely rare in bird genomes and they did not show any activity during last 30 MY, yet they are present in the GRCs of all four examined species, being apparently inherited from the common GRC ancestor. This provides a further evidence for the GRCs to form in the songbird genome rather than in the older avian ancestors because GRCs had a chance to accumulate LTRs while SINEs was likely transferred from the somatic genome copies and amplified in GRCs.

**Table 2.** Fractions and families of interspersed repetitive elements in the GRC sequences and the zebra finch genome.

|  | Zebra finch<br>genome | Zebra finch<br>GRC | Bengalese<br>finch GRC | Eurasian<br>siskin GRC | Pale martin<br>GRC |
| --- | --- | --- | --- | --- | --- |
| Retroelements | 6.97% | 7.62% | 4.69% | 3.99% | 5.75% |
| SINEs | 0.076% | 0.01% | 0.003% | 0.046% | 0.024% |
| LINEs | 3.45% | 0.50% | 0.29% | 1.27% | 1.75% |
| CR1 | 3.40% | 0.49% | 0.28% | 1.27% | 1.75% |
| Other LINEs | 0.05% | 0.01% | 0.01% | 0.00% | 0.00% |
| LTR elements | 3.44% | 7.12% | 4.40% | 2.67% | 3.98% |
| DNA transposons | 0.20% | 0.01% | 0.01% | 0.01% | 0.01% |
| Unclassified: | 0.04% | 0.00% | 0.00% | 0.00% | 0.01% |
| Total interspersed repeats | 7.20% | 7.64% | 4.70% | 4.00% | 5.77% |
| Small RNA | 0.02% | 0.01% | 0.01% | 0.01% | 0.00% |
| Satellites | 0.01% | 0.00% | 0.00% | 0.00% | 0.00% |
| Simple repeats | 1.02% | 2.59% | 2.72% | 2.48% | 2.99% |
| Low complexity | 0.24% | 2.64% | 2.87% | 2.70% | 2.71% |
| Bases masked | 8.48% | 12.88% | 10.29% | 9.19% | 11.46% |

Recently, using transcriptome sequencing of germ cells, an *α-SNAP* gene specific to zebra finch GRC and absent in the somatic genome has been found^5^. However, it is not clear whether the transcription of the *α-SNAP* in the testes and ovaries of zebra finch leads to a functional protein. To examine the general pattern of GRC functioning in oogenesis we analyzed the lampbrush GRCs isolated from the zebra finch oocytes at previtellogenic growth phase. The lampbrush GRC exhibited a typical chromomere-loop pattern, with several pairs of transcriptionally active lateral loops extending from each chromomere, except those located in a prominent DAPI-positive region. Antibodies against the RNA-polymerase II labeled the whole GRC except for this region (Supplementary Fig. 4). Thus, the lampbrush GRCs displays the pattern of transcription typical to all lampbrush chromosomes, which generally do not produce RNA of protein-coding sequences^23^. Therefore, the results of transcriptome analysis of ovaries should be considered with caution in the search for functional GRC genes.

Thus, we discovered that GRC is present in all songbirds investigated and absent in all other birds. We found that GRCs of different songbird species vary drastically in sizes and show low homology between each other. They contain various highly duplicated parts of the somatic genome as well as many repetitive sequences. The spectrum of transposable elements in sequenced GRC libraries suggests that GRC was formed in the songbird lineage rather than in the ancestral avian lineage. We propose that GRC formed as an additional microchromosome in the ancestral songbird genome, which then accumulated additional sequences in some of the species while in others it stayed as a microchromosome (Fig. 3). Aneuploidy for microchromosome could not affect seriously fitness of its carriers. If these proto-GRCs contained extra copies of genes controlling gametogenesis they could even be beneficial. Elimination of such genes (or most of their copies) from the somatic genome would have given the songbirds a substantial economy of the somatic cell size, thus improving their metabolic efficiency, and would be strongly supported by natural selection for a small genome size^2^. The germline restriction of this chromosome would relax natural selection for the functional integrity of its genetic content and make the GRC an easy vehicle for retrotransposons, other selfish genetic elements and amplified copies of unique fragments of the somatic genome. Crossing over suppression along the GRCs (except their termini in female meiosis) could facilitate their divergence and degradation of their original genetic content via the Muller’s ratchet mechanism^24^. These factors might lead to a rapid and massive loss of homology between the descendants of the ancestral GRC. Since GRC persists in the germ line of every songbird studied, it should contain functional genetic element(s) indispensable for the gametogenesis, which are yet to be discovered. Thus, GRCs of songbirds provide a fascinating model for studies in various fields of evolutionary and cell biology.

**Fig. 3.**
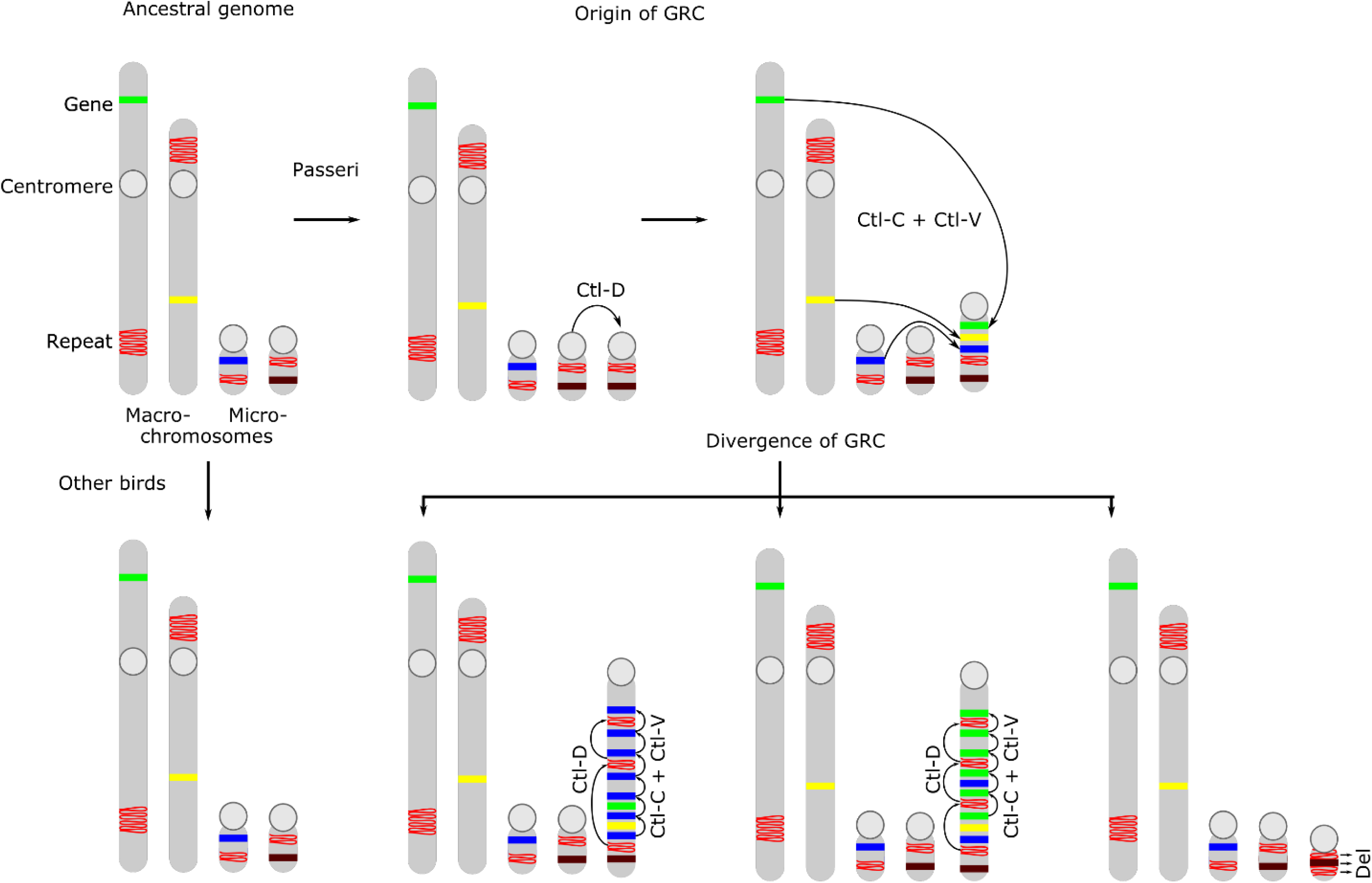
Scenario of GRC origin and evolution.

## Acknowledgments

We thank M.I. Rodionova for the help in chromosome preparation, A. Maslov, D. Taranenko, I. Korobitsyn, M. Scherbakova for the help in bird collecting and the Microscopic Center of the Siberian Department of the Russian Academy of Sciences for granting access to microscopic equipment.

This work was supported by the Russian Foundation for Basic Research (Grant # 18-04-00924) and the Ministry of Science and High Education via the Institute of Cytology and Genetics (Grant # 0324-2018-0019). The funding bodies play no role in the design of the study and collection, analysis, and interpretation of data and in writing the manuscript.

## Author contributions

A.A.T., L.P.M. and I.E.P. prepared and analyzed meiotic and somatic chromosome preparations, K.S.Z, T.V.K. and N.B.R. prepared GRC DNA probes and libraries and performed FISH experiments, E.A.K and E.P.S. collected birds and isolated samples for the analysis, V.A.V., A.F.S. and S.A.G. prepared and analyzed lampbrush chromosome preparations, A.A.T., L.P.M. and D.M.L. analyzed sequence data, A.A.T.,S.A.G., D.M.L., N.B.R. and P.M.B. wrote the paper, A.A.T. and P.M.B designed and oversaw the study.

## METHODS

### Experimental model and subject details

Adult male pale martin, great tit, barn swallow, European pied flycatcher, Blyth’s reed warbler and black tern were captured at the beginning of breeding season. Nestling female sand martin, pale martin, barn swallow, great tit, and European pied flycatcher ∼3-6 days after hatching were collected from the nests.

Adult male zebra finch, Lady Gouldian finch, Bengalese finch, Eurasian siskin, European goldfinch, Eurasian skylark, pine bunting, Eurasian bullfinch and budgerigar were purchased from a commercial breeder. Sexually mature zebra finch females were provided by the Leningrad Zoo (Saint Petersburg, Russia). An adult male rook with fatal accident trauma was provided by the Bird Rehabilitation Centre of Novosibirsk and euthanized in our laboratory.

Capture, handling and euthanasia of the birds followed protocols approved by the Animal Care and Use Committee of the Institute of Cytology and Genetics SB RAS (protocol #35 from 26.10.2016) and by the Saint Petersburg State University Ethics Committee (statement # 131-03-2). Experiments described in this manuscript were carried out in accordance with the approved national guidelines for the care and use of animals. No additional permits are required for research on non-listed species in Russia.

### Mitotic metaphase chromosomes

Mitotic chromosome preparations were obtained from short-term bone marrow cell cultures incubated for 2 h at 37°C in culture Dulbecco’s Modified Eagle’s medium with UltraGlutamine with 10 µg/ml colchicine. Hypotonic treatment was performed with 0.56% KCl solution for 15 min at 37°C and followed by centrifugation for 5 min at 500×*g*. Fresh cold fixative solution (methanol : *glacial* acetic acid, 3:1) was changed three times. Cell suspension was dropped on the cold wet slides slides (76 mm x 26 mm, 1 mm thick). The slides were dried for 2 hours at 65°C and stained for 4 min with 1 μg/ml solution of DAPI in 2×SSC. Then slides were washed in deionized water, dried at room temperature and mounted in Vectashield antifade mounting medium (Vector Laboratories) to reduce fluorescence fading.

### Spermatocyte spreading and immunostaining

Chromosome spreads for SC analysis were prepared from spermatocytes or juvenile oocytes according to Peters et al^25^. Immunostaining was performed according to the protocol described by Anderson et al.^26^ using rabbit polyclonal anti-SYCP3 (1:500; Abcam), mouse monoclonal anti-MLH1 (1:50; Abcam), and human anticentromere (ACA) (1:100; Antibodies Inc) primary antibodies. The secondary antibodies used were Cy3-conjugated goat anti-rabbit (1:500; Jackson ImmunoResearch), FITC-conjugated goat anti-mouse (1:50; Jackson ImmunoResearch), and AMCA-conjugated donkey anti-human (1:100; Jackson ImmunoResearch). Antibodies were diluted in PBT (3 % bovine serum albumin and 0.05 % Tween 20 in phosphate-buffered saline). A solution of 10% PBT was used for blocking. Primary antibody incubations were performed overnight in a humid chamber at 37°C; and secondary antibody incubations, for 1 h at 37°C. Slides were mounted in Vectashield antifade mounting medium (Vector Laboratories) to reduce fluorescence fading.

### Lampbrush chromosome preparations

Zebra finch lampbrush chromosomes were manually dissected from previtellogenic or early vitellogenic oocytes using the standard avian lampbrush technique described in Saifitdinova et al^27^. After centrifugation, preparations were fixed in 2% paraformaldehyde, then in 50% and in 70% ethanol, air-dried and kept at room temperature until used for FISH. For immunostaining experiments lampbrush chromosome preparations were kept in 70% ethanol at +4°C.

### The preparation of the hybridization probe and FISH

In order to generate DNA probe for GRC of the pale martin, zebra finch, Bengalese finch and Eurasian siskin testicular cells of adult males were treated with hypotonic solution (0.88% KCl) at 37° for 3h and then with Carnoy’s solution (methanol : *glacial* acetic acid, 3:1). The cell suspension was dropped onto clean cold wet cover slips (60 mm x 24 mm, 0.17 mm thick), dried, and stained with 0.1% Giemsa solution (Sigma) for 3-5 min at room temperature. GRCs were identified as positive round bodies located near the spermatocytes I. Microdissection of GRC and amplification of DNA isolated from this chromosome by GenomePlex Single Cell Whole Genome Amplification Kit (WGA4) (Sigma-Aldrich)^28^. Microdissected DNA probes were generated from 15 copies of GRC for each studied species. The obtained PCR products were labeled with Flu-dUTP (Genetyx, Novosibirsk) in additional PCR cycles or with biotin-11-dUTP (Sileks, Moscow, Russia).

FISH experiments with DNA probes on SC spreads of the studied avian species were performed as described earlier^29^ with salmon sperm DNA (Ambion, USA) as a DNA carrier. In case of suppression FISH, Cot-1 DNA (DNA enriched for repetitive DNA sequences) was added to DNA probe to suppress the repetitive DNA hybridization. Chromosomes were counterstained with DAPI dissolved in Vectashield antifade solution (Vector Laboratories, USA). Zebra finch GRC at the lampbrush stage was identified by FISH using biotin-labelled zebra finch microdissected probe with a 50-fold excess of *E. coli* tRNA as a carrier. FISH was performed according to the DNA/DNA+RNA hybridisation protocol omitting any chromosome pre-treatment, as described previously^30^. To detect biotin-labelled probe, we used avidin-Alexa488 and biotinylated goat antibody against avidin (both from Thermo Fisher Scientific, USA). Lampbrush chromosomes were counterstained with DAPI in antifade solution, containing 50% glycerol.

### Immunostaining of the zebra finch lampbrush chromosomes

Immunostaining was carried out with mouse antibodies V22 (kindly donated by U. Scheer) against the phosphorylated C-terminal domain (CTD) of RNA polymerase II. Lampbrush chromosome spreads, fixed in 2% paraformaldehyde, were blocked in 0.5% blocking reagent (Sigma-Aldrich, USA) in PBS for 1 h at +37°C. Then preparations were incubated with primary antibodies, diluted 1:200, overnight at room temperature. Slides were washed in PBS with 0.05% Tween-20 and incubated with Alexa-488-conjugated goat anti-mouse IgG+IgM secondary antibody (Jackson ImmunoResearch Lab). After washing in PBS+0.05% Tween-20, slides were mounted in antifade solution containing DAPI.

### Microscopic analysis

Images of fluorescently stained metaphase chromosomes and/or SC spreads were captured using a CCD-camera installed on an Axioplan 2 compound microscope (Carl Zeiss, Germany) equipped with filtercubes #49, #10, and #15 (ZEISS, Germany) using ISIS4 (METASystems GmbH, Germany) at the Center for Microscopic Analysis of Biological Objects of SB RAS (Novosibirsk, Russia). For further image analysis of we used Corel PaintShop Pro X6 (Corel). The location of each imaged immunolabeled spread was recorded so that it could be relocated on the slide after FISH. Zebra finch lampbrush chromosome preparations were examined using a Leica DM4000B fluorescence microscope installed at the “Chromas” Resource Centre, Saint-Petersburg State University Scientific Park (Saint-Petersburg, Russia). The microscope was equipped with a black and white DFC350FX camera and filters A and I3. LAS AF (Leica) software was used to capture and process color images; Adobe Photoshop CS5 (Adobe Systems) was used for figure assembling.

### Preparation of amplified DNA and library construction

DNA amplification of microdissected GRC chromosomal material was performed by GenomePlex Single Cell Whole Genome Amplification kit (WGA4) (Sigma-Aldrich) according to the manufacturer protocol. DNA library for NGS sequencing was prepared based on the microdissected GRC DNA libraries using NEBNext Ultra DNA Library Prep kit (New England Biolabs).

### High throughput sequencing and error correction

NEBNext Ultra library was sequenced on an Illumina NextSeq 5500 system with single-end reads at the “Genomics” core facility of the ICG SB RAS (Novosibirsk, Russia). Read lengths were 150 bp, the total number of reads obtained were 1,730,845, 1,596,722, 2,821,862 and 1,265,105 for zebra finch, Bengalese finch, Eurasian siskin and pale martin GRC correspondingly. DNA data were quality assessed using FastQC^31^ and quality trimmed using Trimmomatic^32^.

### Estimating the homeology to somatic genome and repeat content

The reads from zebra finch, Bengalese finch, Eurasian siskin, and pale martin GRC sequences were aligned to the assembly of the zebra finch genome (Taeniopygia_guttata-3.2.4^14^) using BLAT^15^. A custom python script was used to estimate the coverage of the zebra finch genome in 10 kb windows^16^. Overlapping of the regions with high coverage with zebra finch Ensembl gene predictions and non-zebra finch RefSeq^17^ genes was revealed with Ensembl genome browser. The repeat content of the GRC libraries and zebra finch genome was assessed with RepeatMasker^19^ by using avian RepBase database^20^.

### Mapping of the GCR sequence aligned to unique region of zebra finch genome

To confirm homology between GRC sequences and a zebra finch genome we designed a DNA probe based on the 1.5 kb long sequence from the region of TGU5:31550464-31551880. Region of interest was amplified using zebra finch DNA isolated from somatic tissues and primers designed with Primer-BLAST tool using 35 PCR cycles. The obtained PCR products were labeled with Flu-dUTP (BIOSAN, Novosibirsk) in 25 additional PCR cycles. FISH with the probes on SC spreads were performed according to a standard protocol^33^.

### Quantification and statistical analysis

The length of the SC of each chromosome arm was measured in micrometers and the positions of centromeres were recorded using MicroMeasure 3.3^34^. We identified individual SCs by their relative lengths and centromeric indexes.

## Supplemental information

**Supplementary Figure1.**
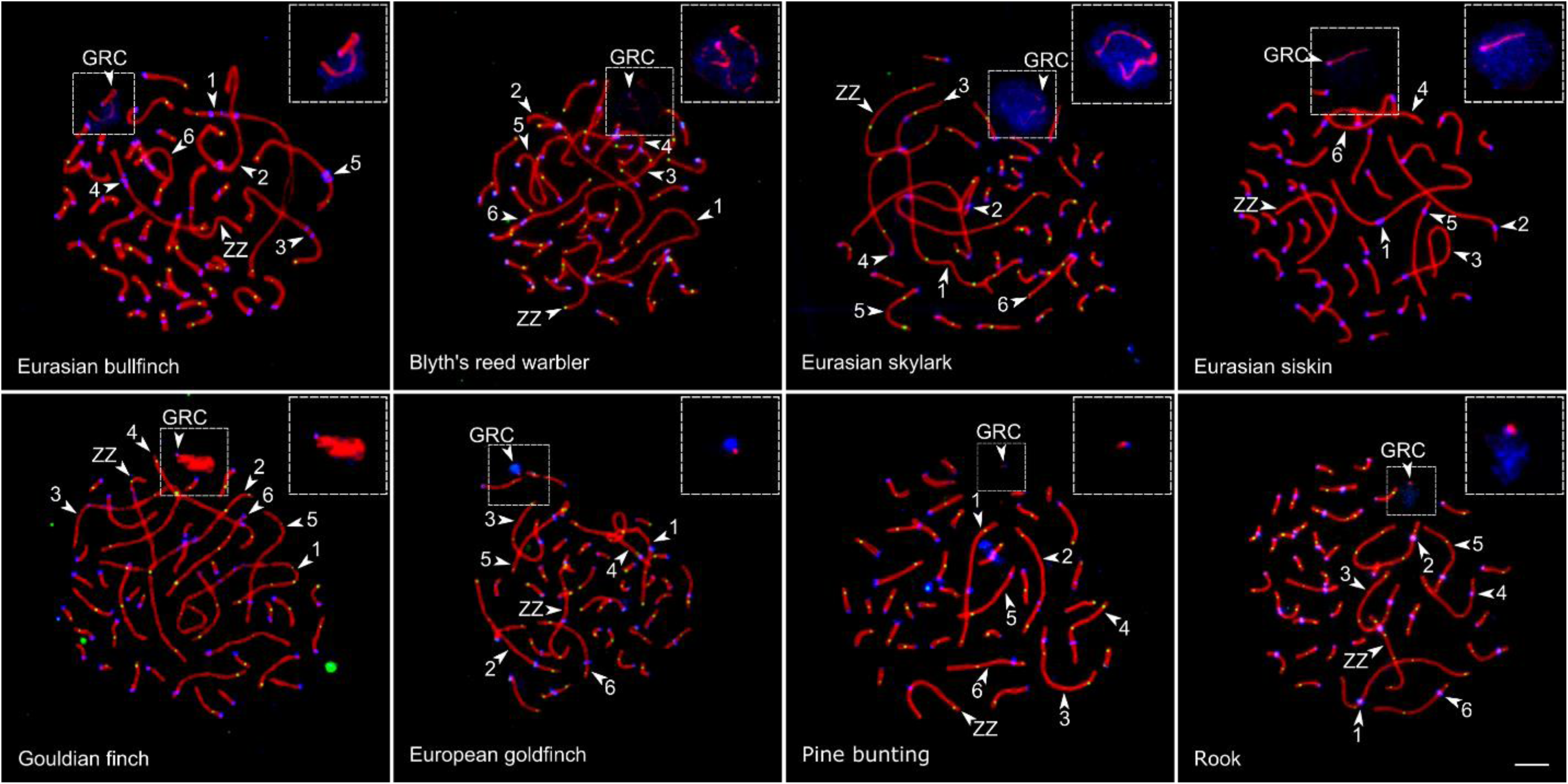
Discovery of GRCs in bird species. SC spreads of the spermatocytes of eight bird species immunolabelled with antibodies against SYCP3 (red), centromere proteins (blue) and MLH1 (green). Arrowheads point to the largest chromosomes ordered according to their size, ZZ (identified by its size and the arm ratio), bivalents, and GRCs. GRC is represented as an acrocentric macrochromosome (Eurasian bullfinch, Blyth’s reed warbler, Eurasian skylark, Eurasian siskin and Gouldian finch) or microchromosome (European goldfinch, pine bunting and rook). In most spermatocytes of all species, GRC SC was surrounded by a cloud of chromatin labelled with anticentromere antibodies. Bar – 5 µm.

**Supplementary Figure2.**
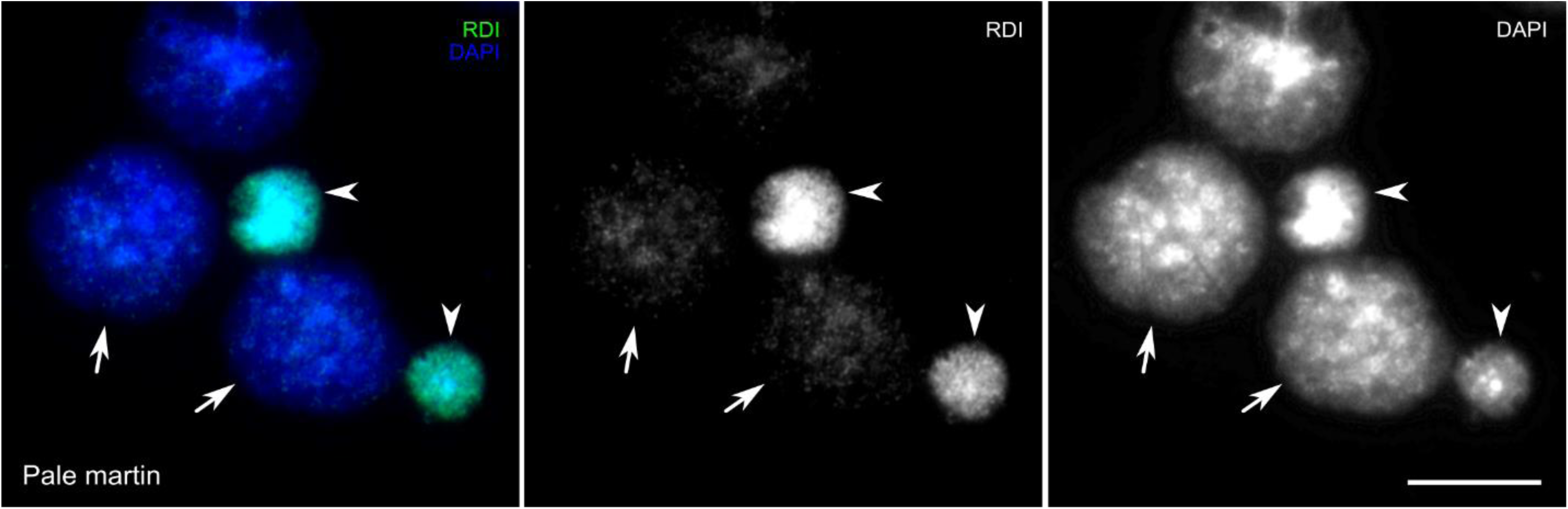
Visualization of eliminated GRC. Spread of the pale martin germ cells after FISH with pale martin GRC probe (green) and DAPI staining (blue). The round dense bodies (pointed by arrowheads) containing GRC eliminated from the spermatocytes (arrows) were used for microdissection and preparation of DNA libraries. Bar – 10 µm.

**Supplementary Figure3.**
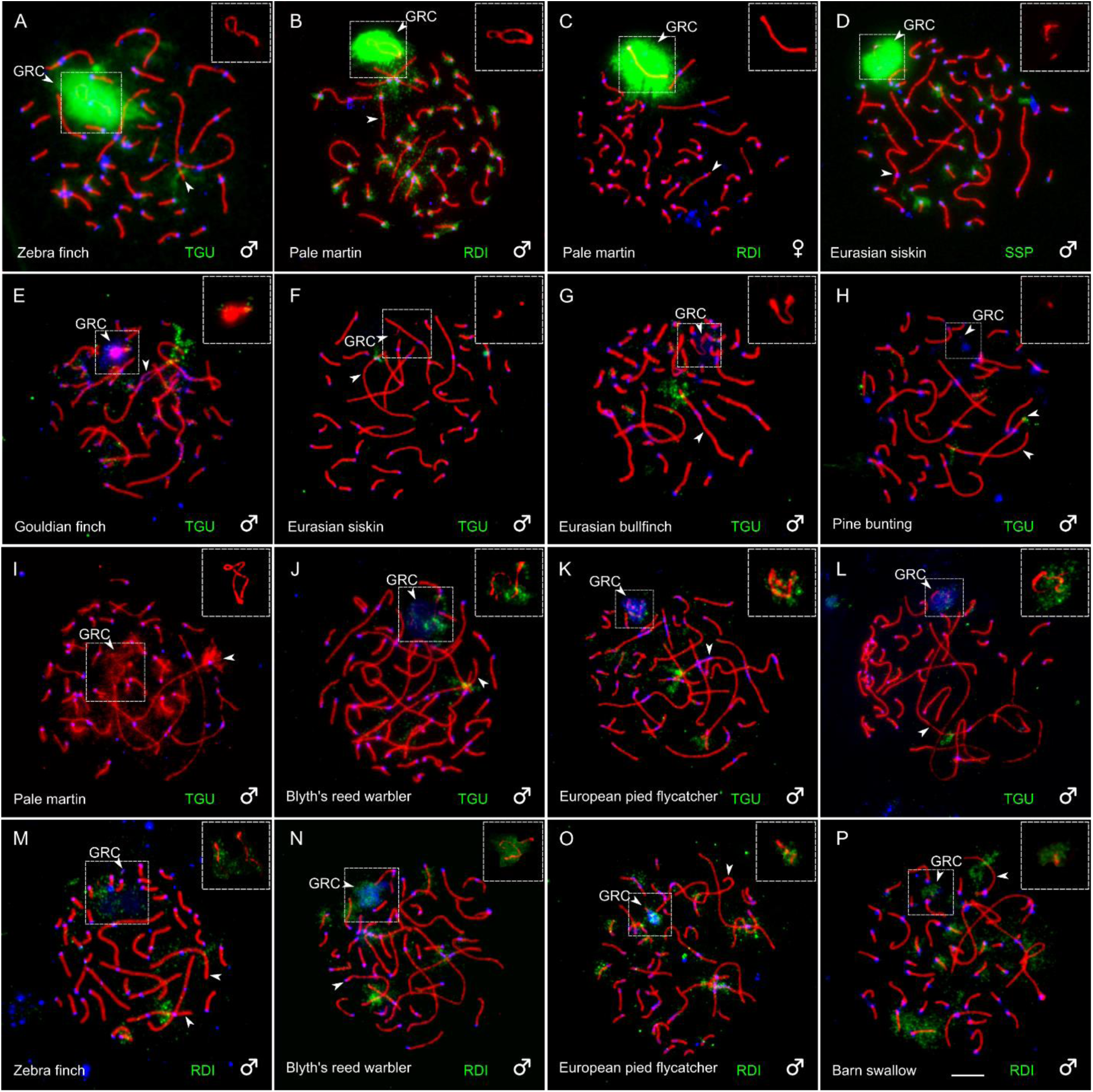
FISH with GRC-specific microdissected DNA probes. Reverse (A-D) and cross-species (E-P) FISH of DNA probes (green) derived from GRC of zebra finch (TGU), pale martin (RDI), and Eurasian siskin (SSP) with SC spreads, immunolabelled with antibodies against SYCP3 (red). Centromeres are labeled with anticentromere antibodies (blue). Inserts show GRCs. In reverse FISH, GRC probes produce strong specific signal at the GRC. The zebra finch GRC probe produces specific signal at the short arm of SC 3 and a weak dispersed signal at some macro- and microchromosomes. The pale martin GRC probe paints the pericentromeric regions of all chromosomes of the somatic chromosome set. The Eurasian siskin GRC probe gives specific signals at the long arm of SC 3, some pericentromeric and other regions. In cross-species FISH, the zebra finch GRC probe produces almost no signal at GRC of Gouldian finch (E), Eurasian siskin (F), Eurasian bullfinch (G) and pine bunting (H), and slightly paints GRCs of pale martin (I), Blyth’s reed warbler (J), European pied flycatcher (K), and Eurasian skylark (L). In all species, it produces a specific signal at the short arm of SC 3. The pale martin GRC probe produces a weak specific signal at GRC of zebra finch (M), Blyth’s reed warbler (N), European pied flycatcher (O), and barn swallow (P). In all species, this probe produces signal at SC4 and 5. In Blyth’s reed warbler (N), European pied flycatcher (O), and barn swallow (P) it also paints some microchromosomes. Bar – 5 µm.

**Supplementary Figure4.**
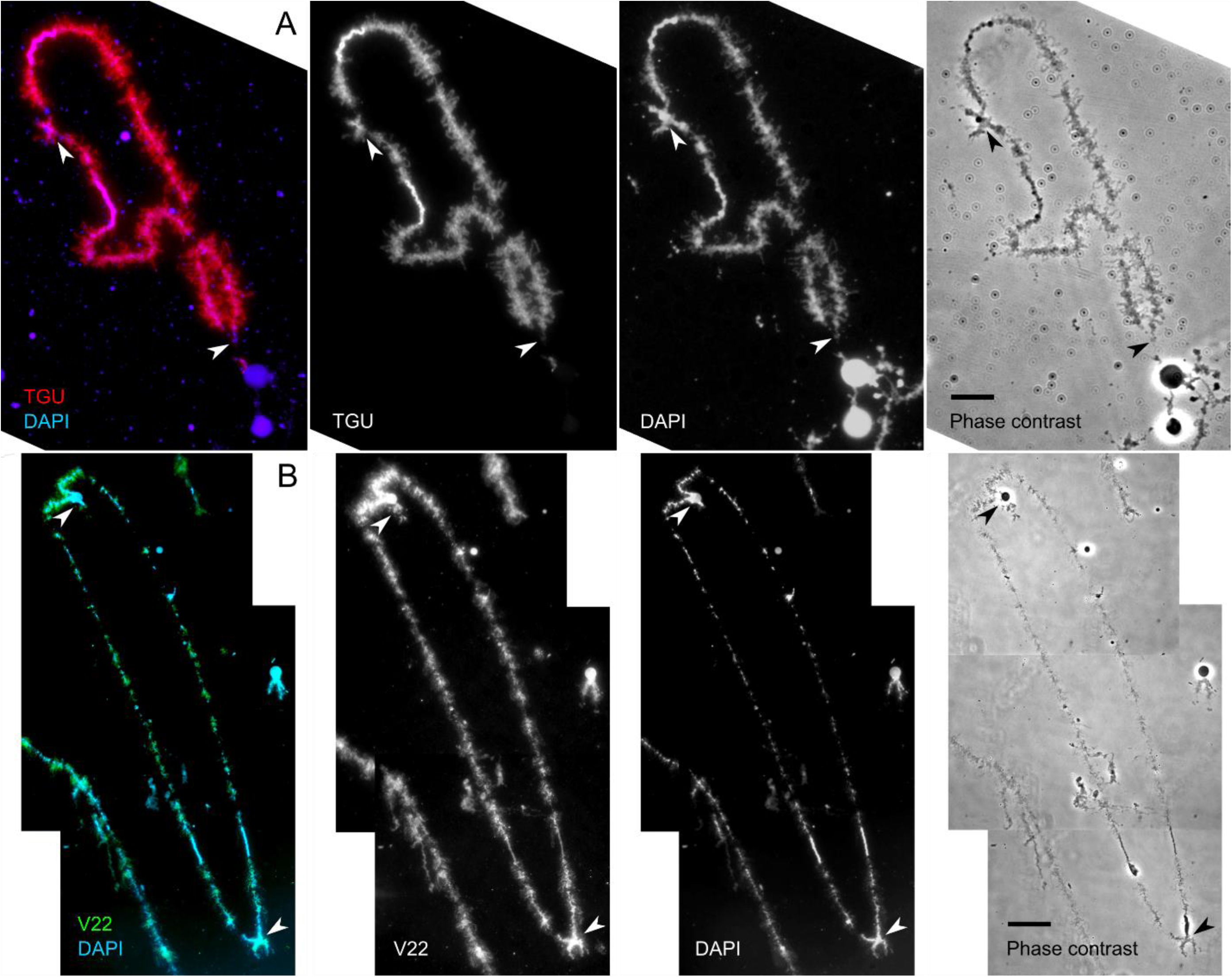
Zebra finch lampbrush GRC. (A) Identification of lampbrush GRC using FISH with zebra finch whole chromosome microdissected probe (red). (B) Immunodetection of the phosphorylated form of RNA polymerase II with V22 antibody. The axes of lateral loops are immunolabelled (green). (A, B) DAPI stained chromosomes (blue). Arrows indicate chiasmata. Bars - 20 μm

